# Using TMS-EEG to study the intricate interplay between GABAergic inhibition and glutamatergic excitation during reactive and proactive motor inhibition

**DOI:** 10.1101/2024.09.05.610957

**Authors:** Tingting Zhu, Chenglong Cao, James Coxon, Alexander T. Sack, Inge Leunissen

## Abstract

Transcranial magnetic stimulation (TMS) studies have provided insight into the neurotransmission underlying motor inhibition but are limited to the primary motor cortex (M1). To investigate γ-aminobutyric acid (GABA) and glutamatergic neurotransmission across the broader motor inhibition network, we combined TMS with electroencephalography (EEG). The N45 and N100 components of the TMS-evoked potentials (TEPs) have been liked to GABAa and GABAb signaling, respectively, whereas the N15, P30 and P60 TEPs are believed to reflect glutamatergic neurotransmission. This study investigates how these components are modulated during reactive and proactive motor inhibition. Twenty-four healthy participants completed two TMS-EEG sessions targeting either the left M1 or the pre-supplementary motor area (preSMA) at rest and during a stop-signal task. We compared TEP amplitudes across TMS conditions (active/sham) and trial types and assessed their relationship with task performance. Participants with a higher N45 amplitude over M1 at rest demonstrated a faster stop-signal reaction time. Active TMS during successful motor inhibition evoked higher N15 amplitudes over preSMA than during uncertain go trials, suggesting that an early excitatory signal from preSMA is crucial for reactive inhibition. In contrast, proactive inhibition was linked to lower P30 amplitudes in M1, possibly reflecting reduced motor facilitation. These findings provide novel empirical insights into the neurophysiological mechanisms underpinning motor inhibition, emphasizing the importance of a coordinated balance between inhibitory processes in M1 and early glutamatergic excitatory signals from preSMA for effective inhibition.

## Introduction

Motor inhibition, an essential function in daily life, refers to the ability to suppress initiated movements, particularly when execution is undesirable (Duque et al., 2017; Verbruggen et al., 2019). Two variants of motor inhibition can be distinguished, reactive and proactive inhibition (Aron, 2011). While reactive inhibition represents motor suppression in reaction to an unexpected stop signal during an ongoing movement, proactive inhibition refers to a preparatory process that facilitates movement cancellation when one might be required to stop soon. The putative neural underpinnings involve complex interactions within a fronto-basal ganglia network, wherein the right inferior frontal cortex and pre-supplementary motor area (preSMA) instigate inhibition of thalamo-cortical output via the indirect and hyperdirect cortico-basal ganglia pathways (Jahanshahi et al., 2015).

Given the prominent role of the primary motor cortex (M1) in shaping descending motor output, M1 is likely the site where movement preparation and cancellation meet (Stinear et al., 2009). Single-pulse transcranial magnetic stimulation (spTMS) studies revealed that cortical excitability as measured by motor evoked potentials (MEP) is reduced in M1 during movement cancellation (Badry et al., 2009; Coxon et al., 2006; MacDonald et al., 2014; van den Wildenberg et al., 2010), but this reduction of MEPs could be the consequence of either an active inhibition process or a reduction of excitation. Subsequent paired-pulse TMS (ppTMS) studies demonstrated that motor inhibition indeed coincided with an increase in short intracortical inhibition (SICI) (Coxon et al., 2006; Hermans et al., 2018), and that the degree of this increase relates to the speed of the inhibition process (Chowdhury et al., 2020). This supports that movement cancellation requires an active inhibition process at the level of M1 brought about by an increase in GABAa signaling.

The GABAa receptor is a fast-acting ligand-gated chloride channel that quickly inhibits signals via postsynaptic membrane hyperpolarization, while the GABAb receptor produces prolonged inhibitory effects by G-protein-coupled modulation of intracellular signaling pathways (Terunuma, 2018). It is tempting to map this distinction between the receptor types to the fast reactive inhibition versus prolonged proactive inhibition. GABAb-mediated intracortical inhibition in M1 can also be assessed with ppTMS as long intracortical inhibition (LICI). LICI is indeed increased when a stop signal might appear (Cowie et al., 2016). However, there are also some indications that GABAb signaling is relevant for reactive inhibition, with the cortical silent period prolonged during successful inhibition (van den Wildenberg et al., 2010). When measured at rest, the silent period, LICI, and SICI all relate to inhibitory task performance (He et al., 2024; Loomes et al., 2023; Paci et al., 2021). Measuring LICI during the inhibition process is difficult due to the long interval between the conditioning and test pulses. A potential solution to achieve a more comprehensive understanding of GABAergic neurotransmission during motor inhibition, is to combine TMS with electroencephalography (EEG).

The TMS-evoked potential (TEP) in the EEG signal (Chung et al., 2015; Gordon et al., 2023b; Rogasch et al., 2013; Rogasch & Fitzgerald, 2013; Tremblay et al., 2019) contains a series of peaks, reflecting the excitability and reactivity of the underlying cortical networks (Darmani & Ziemann, 2019; Farzan, 2024). Whereas the N15, P30 and P60 peaks are believed to reflect excitation (Esser et al., 2006; Maki & Ilmoniemi, 2010; Rogasch et al., 2013), the N45 and N100 have been linked to GABAa dependent and GABAb receptor-mediated neurotransmission, respectively (Darmani et al., 2016; Premoli et al., 2014a; Rogasch et al., 2013). An earlier spTMS-EEG study found increased N100 amplitude during successful reactive inhibition (Yamanaka et al., 2013), suggesting GABAb involvement in motor inhibition. Yet, methodological limitations such as a small sample size (N = 6) and lack of a sham condition limit the generalizability of these results.

A major advantage of TMS-EEG is that the assessment of intracortical inhibition and facilitation is no longer limited to M1. Therefore, our study aims to investigate changes in inhibition and excitation during reactive and proactive inhibition by applying a spTMS-EEG protocol targeting either preSMA or left M1 in separate sessions. For M1, we hypothesized that the amplitude of the N45 peak (reflecting GABAa) will be amplified specifically during successful stopping (i.e., reactive inhibition), whereas the N100 peak may exhibit differences already at the start of the trial, indicating the involvement of the slower GABAb system in proactive inhibitory control. The preSMA sends an excitatory signal to the subthalamic nucleus (STN) and striatum when there is a need for inhibition which might result in higher N15, P30 or P60 amplitudes (reflecting glutamate) during reactive and/or proactive inhibition. Understanding the intricate interplay between GABAergic inhibition and glutamatergic excitation can contribute to a better understanding of the mechanisms of motor inhibition and provide valuable insights into potential treatments for neurological and psychiatric disorders with inhibitory dysfunction.

## Methods

### Participants

Twenty-five healthy participants were recruited for this study (11 males, age range: 19-31, mean age: 25.56 ± 4.35). The inclusion criteria were: (i) right-handed (mean laterality quotient: 97.5 ± 5.95, as assessed by the Oldfield handedness inventory (Oldfield, 1971)); (ii) no history of neurological disorders (such as epilepsy, seizure or stroke) or neuropsychiatric disorders; (iii) no history of alcohol or other drug abuse/dependence; and (iv) no TMS and MRI contraindications, such as the metal implants (Rossi et al., 2009). All participants gave their written informed consent and were remunerated for their participation. The experimental procedure was approved by the local ethics committee.

### Experimental design

The study consisted of three separate sessions, conducted on different days. In the first session, we obtained a structural MRI, and a 12-min stop-signal task (SST) assessment was carried out to measure participants’ baseline inhibitory performance. During the second and third sessions (minimally 48h apart, range = 48 - 435 h, mean = 213 ± 123 h), participants underwent TMS-EEG targeting either preSMA or M1. TMS was applied at rest (2 blocks: 1 active stimulation and 1 sham) and during SST performance (4 blocks: 2 active stimulations, 2 sham). The order of the active and sham stimulation was counterbalanced over participants but kept the same for both targets within subjects.

### MRI data acquisition

A structural MRI was acquired for every participant to navigate the TMS coil. MR scanning was performed on a Siemens Prisma 3T scanner with the following parameters: 192 sagittal slices, TR = 2300ms, TE = 2.98ms, FOV = 256 × 256 mm^2^, voxel size = 1 mm×1mm×1mm, flip angle = 9°.

### Stop-signal Task paradigm

An anticipatory response version of the SST, programmed in LabView (2020), was used to assess motor inhibition (Figure 1). Participants were seated ∼1m from a 240Hz screen. The display consisted of a vertical bar that ascended at a constant speed, reaching the top in 1000ms. Participants were instructed to press a button (right index finger) to halt the bar at the target line (800ms) as accurately as possible during go trials. In 25% of trials, the bar unexpectedly stopped before the target line. In these stop trials participants were expected to withhold their response. During the baseline assessment, a staircasing algorithm kept P(inhibit) ≈ 50%. The initial stop signal delay (SSD), i.e. the time into the trial at which the filling bar stops was set 200ms from target and adjusted in steps of 25ms. We instructed participants to respond as accurately as possible on go trials, and that it would not be possible to withhold the button press on all stop trials. Participants performed two practice blocks: 20 go trials, then 20 trials comprising stop and go trials. The task commenced with 40 go only trials (light blue bar, certain go trials), followed by 200 trials (dark blue bar) comprising 150 go trials (uncertain go trials) and 50 stop trials randomly intermixed. The inter-trial interval (ITI) ranged from 3 to 4 seconds.

**Figure 1.**
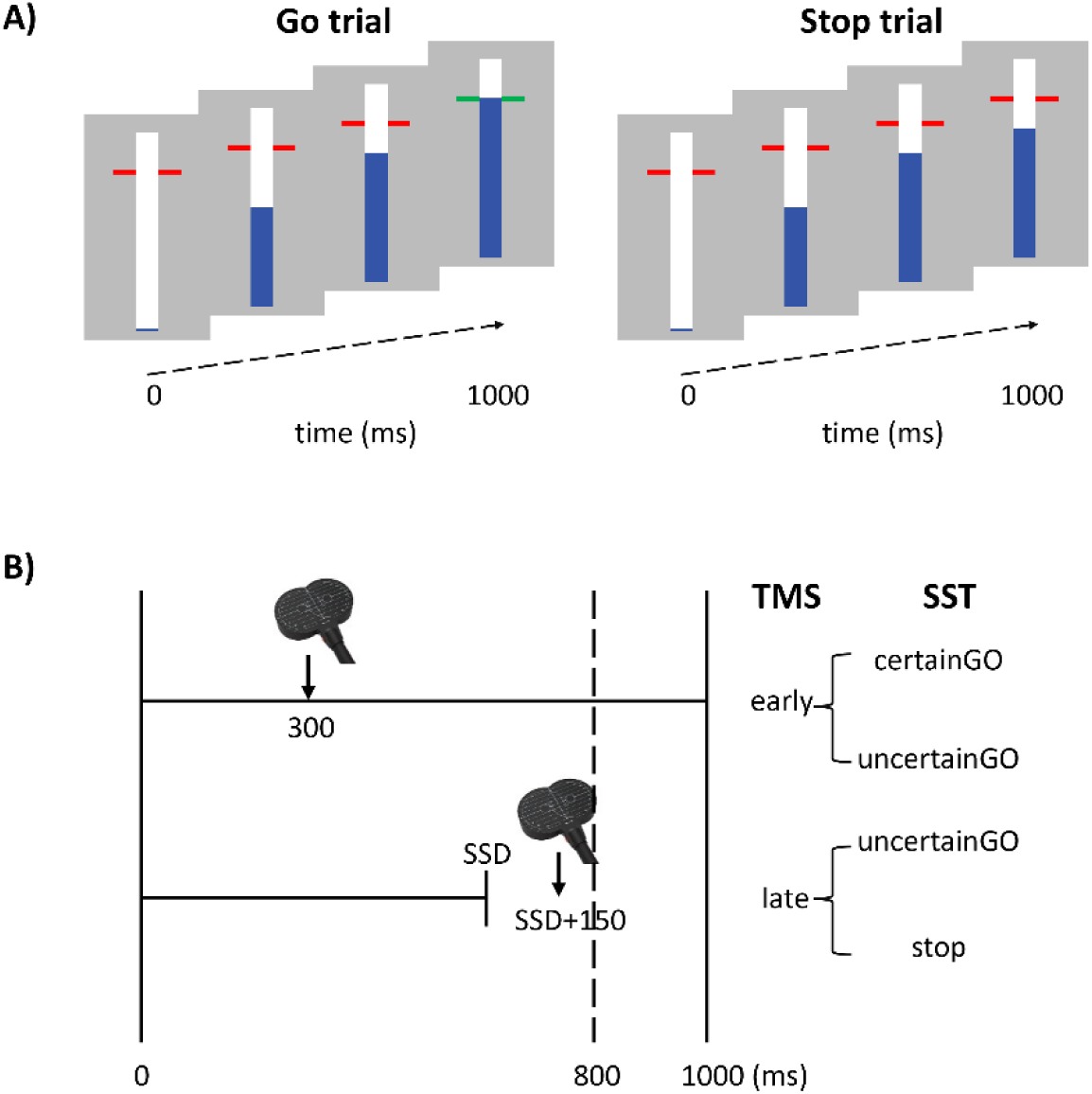
Stop-signal task paradigm with timing of TMS and conditions of interest. (**A)** The anticipatory version of the SST display with mixed go and stop trials. Participants were asked to halt an ascending bar at the target line during go trials. Feedback was given based on response timing (green <20ms, yellow <40ms, orange <60ms, or red >60ms from target). Stop trials required response, withholding when the bar stopped earlier than the target line, no feedback was provided on stopping accuracy. (**B**) Experimental design indicating the timing of TMS pulse on SST trials. The arrow presents the time when TMS pulse was given. The early TMS was administered 300ms after trial onset, while late TMS pulses occurred 150ms after the stop signal presentation. The early TMS condition occurred on both certain and uncertain go trials, but the late TMS occurred on uncertain go and stop trials. SSD = stop signal delay.

### TMS-EEG set-up

EEG was recorded with 64 low-profile electrodes (Braincap-TMS, Easycap GmbH) linked to a BrainAmp DC Amplifier (Brain Products, GmbH) with ground electrode located at FPz and the online reference at TP10. Electrooculography signals capturing horizontal and vertical eye movements, as well as the electromyography (EMG) signal from the first dorsal interosseous (FDI) muscle was acquired using an additional BrainAmp ExG amplifier (Brain Products, GmbH). For the EMG disposable adhesive electrodes (∅20 mm) we placed in bipolar belly-tendon electrode montage (GND: processus styloideus ulnaris). Data was recorded from DC with a lowpass filter of 1 kHz and digitized at a sampling rate of 5 kHz. For EEG electrodes impedances were kept below 5 kΩ and the electrode leads were oriented perpendicular to the coil handle. Polymeric foam spacers (∼0.5 cm high) were placed on the EEG cap to avoid direct contact of the TMS coil with EEG electrodes and to minimize bone conduction of the TMS click (Rydalch et al., 2019).

Biphasic TMS pulses were delivered using a MagPro-X100 Stimulator with a 70 mm figure-of-eight coil in a P→A current direction (MC-B70, MagVenture A/S, Farum, Denmark). The FDI motor hotspot located at which five consecutive TMS pulses produced the highest mean motor evoked potential (MEP), coil 45° to the midsagittal plane, handle pointing posteriorly. The motor hotspot was marked on the cap and captured with a neuronavigation system (Localite TMS Navigator 3.0.63, SW-Version3) to ensure optimal positioning during the M1 session. Next, resting motor threshold (RMT) was determined as the lowest stimulus intensity that elicited five out of ten MEP ≥ 50 µV (peak to peak) (Rossini et al., 1994). PreSMA was defined as MNI coordinate (11, 10, 62) (Power et al., 2011; Wang et al., 2010) and transformed to subject space for neuronavigated targeting with the handle oriented posteriorly (Casula et al., 2022). The stimulation intensity was set at 100% RMT (49.93 ± 5.30 %MSO) and decreased by 1-2% on M1 if interference with index finger extension was observed or self-reported by the participant. During sham stimulation, a 4cm polyethylene foam was attached to the bottom of the coil to prevent any transfer of active TMS pulses to the scalp (Conde et al., 2019).

We took measures to reduce somatosensory and auditory TEP contamination. The sound of the TMS coil was masked by administering adaptive white noise with the same frequency of the TMS click through in-ear headphones fitted inside foam earplugs (E-A-RTONE 3A Earphones) (Conde et al., 2019; Gordon et al., 2021; Massimini et al., 2005). Additionally, we applied Cutaneous Electric Stimulation (CES, DS7A Digitimer Ltd, USA) to mask the sensory experience of active TMS without cortical activation (Conde et al., 2019; Gordon et al., 2021; Lin et al., 2003; Torquati et al., 2002) (see supplementary material for more details). White noise and CES were applied during both active and sham TMS to isolate genuine cortical activation evoked by TMS from potentially confounding sensory effects.

Participants were asked to fill out the visual analog scale (VAS, scale 0 to 10) after each block, referring to the perceived sound and sensation, with 0 representing no perception and 10 maximal perceptions. The VAS included the following items: 1) intensity of auditory sensation; 2) intensity of scalp sensation; 3) area of scalp sensation; 4) pain or discomfort (Gordon et al., 2021). Extra items were applied only after the whole experiment: 1) comfort of the sound; 2) differences between blocks.

### TMS at rest

Participants were asked to sit comfortably, be at rest and avoid any systematic thinking. They were instructed to fixate on a white cross on the screen during TMS at rest (rsTMS) block. Seventy-five consecutive TMS pulses with different ITI (4/4.5/5s) were transferred for each rsTMS block (1 active and 1 sham stimulation), lasting for about 6 mins. The target site was kept the same for rest and task within each session.

### TMS during stop-signal task performance

Per session, the SST with TMS included 160 certain go trials, 480 uncertain go trials and 160 stop trials (total = 800 trials) divided over 4 task blocks (2 active and 2 sham stimulation blocks). The TMS pulse was randomly applied at two different time points: early (i.e., 300ms after trial onset) or late (i.e., participant-specific stop time + 150ms) (Figure 1). The late stimulation point was chosen because active inhibition is expected 150ms after presentation of the stop-signal (Coxon et al., 2006). The participant-specific stop time was determined from the no-TMS session with the formula: 800ms – [mean (GoRT) – mean (stop time)] – 50ms. The average reaction time on go trials (GoRT) was defined as the time between trial onset and pressing the finger from the switch. A GoRT of 800ms reflects a response on the target. By using a participant-specific stop time, the degree of stopping difficulty is comparable across participants. By subtracting 50ms, we ensured >50% successful stop trials and sufficient trials for TEP averaging. To keep the participant alert, a small number of stop trials were catch trials (10 trials that were very easy to inhibit, i.e. extra 100ms; 10 trials that were very hard to inhibit, i.e. less 100ms).

The majority of trials (i.e 70%; 560 out of 800 trials) were stimulation trials (either active or sham TMS). There were 70 TMS pulses for each stimulation condition (active or sham), trial type (certain go, uncertain go, stop trial), and pulse timing (early, late). The trial duration was 3.5, 4 or 4.5s depending on the time point of stimulation in the previous and current trial, such that there was at least 3.5s between two consecutive TMS pulses. Non-stimulated trials comprised 20 certain go, 200 uncertain go and 20 stop trials. Sufficient breaks were provided to participants based on their individual pace, and it took about 1h to complete all four task blocks.

## Data analysis

### Behavioral data analysis

Behavioral analysis was performed on the baseline task session without TMS. GoRT was measured as the time elapsed from the onset of the trial to the participant’s response (button press). GoRTs shorter than 400ms (i.e., early responses) or longer than 1000ms (i.e., no response) were considered errors and removed from the analyses (Leunissen et al., 2016; Leunissen et al., 2022; Leunissen et al., 2017). The average goRT was calculated for certain and uncertain go trials separately. The presence of stop trials in the mixed task blocks introduces uncertainty, as participants are aware that they may have to inhibit their response at any moment. This uncertainty encourages proactive inhibition, where participants slow their responses to prepare for a possible stop signal. It creates a strategic “buffer” time to increase their chance of successfully inhibiting a response on possible stop trials. By contrast, certain go trials (where participants know there are no stop trials) typically produce faster responses since there is no need to anticipate a stop signal (van den Wildenberg et al., 2022). The degree of proactive inhibition was quantified by calculating the difference in RT between uncertain go and certain go trials.

Reactive inhibition refers to the ability to halt a response upon encountering a stop signal. To estimate the speed of this inhibitory process we determined the stop-signal reaction time (SSRT) with the integration method (Verbruggen et al., 2019). More specifically, the number of failed stop trials was divided by the total number of stop trials to get P(respond). Uncertain goRTs were sorted in ascending order and the nth goRT was obtained where n equals the number of go trials multiplied by P(respond). Go omissions were replaced with the maximum RT of 1000ms. The SSRT was estimated by subtracting the average stop time from the nth goRT.

### TMS-EEG data processing

#### Data preprocessing

All data processing was performed in EEGLAB (v2021.1), using the TMS-EEG signal analyzer toolbox (TESA 1.1.1) (Rogasch et al., 2017) on the MATLAB 2021b platform (MathWorks®, Inc., Massachusetts, USA). We followed a processing pipeline based on recommendations from Bertazzoli et al. (Bertazzoli et al., 2021) and Rogasch et al. (Rogasch et al., 2022). Before commencing the data processing, we merged the active and sham TMS blocks for the same target into a single datafile. Bad channels (saturated or noisy) were removed. TMS-EEG data was then epoched around each TMS pulse (−1000 to 1000ms) and baseline-corrected to the pre-TMS pulses period (−200 to -10ms). To remove the large initial artifact of the TMS pulse, we discarded the data 4 to 10ms before and after the TMS pulse, followed by cubic interpolation over a 20ms window.

The data was subsequently downsampled from 5000 to 1000Hz. We then visually inspected and removed bad trials (range = 0-24, mean = 7.3 ± 6.91). An initial independent component analysis (FastICA) (Hyvarinen & Oja, 2000) was applied to remove TMS-related artifacts (muscle, eye blink, decay and recharging artifacts (Rogasch et al., 2017)). After performing bandpass filter from 1 to 100Hz and bandstop filter from 48 to 52Hz (remove non-neural data and line noise), we applied a second round of fastICA to eliminate remaining artifacts, such as eye movements and blinks, muscle and electrode artifacts (Rogasch et al., 2017). Bad channels were then interpolated using spherical interpolation. Lastly, the data was re-referenced using the average of all channels. Finally, the resulting time courses were baseline corrected to the pre-TMS pulse period (−200 to -10ms) again. Due to the event related potential evoked by the onset of the trial, trials with TMS provided at the early timepoint (i.e. 300ms after bar fill onset) were baseline corrected to the -400 to -200ms pre-pulse period. The non-stimulated trials followed the same preprocessing steps with epoching about virtual TMS event markers (omitting steps related to the TMS artifact and the first ICA).

### TEP peak analysis

TEPs were calculated by averaging across trials for each condition. We defined the channel closest to the target site as region of interest to extract the TEP waveform, which was C3 for M1 and FC2 for preSMA. The TEP peak amplitude in a prespecified window (N15 (10-20ms), P30 (20-40ms), N45 (30-60ms), P60 (40-80ms), N100 (80-140ms) and P180 (160-240ms) were then extracted. If no peak was found within the window, the amplitude at the center timepoint was defined as the amplitude for that peak (e.g., the TEP amplitude at 45ms would be defined as N45 if there was no exact N45).

### ERP analysis

Stimulus-locked event-related potentials (ERPs) were created for go trials (time-locked to the average SSD) and stop trials (time-locked to the SSD). We identified the peak of the P2, N2 and P3 by identifying the maximum trough or peak within the window of interest (180-400ms following the stop-signal).

### MEP analysis

EMG data collected during TMS on M1 were processed to compute MEPs. The TMS pulse was first identified, and data were epoched from 1 second before to 1 second after the pulse. Preprocessing included linear detrending and baseline correction, where the baseline was defined as the period from 200 ms to 5 ms prior to the TMS pulse. To assess background EMG activity, the root mean square (RMS) of the EMG signal in the 50 ms preceding the TMS pulse was calculated, and trials with RMS values exceeding 20 µV were excluded to ensure data quality. Additionally, go trials with early or no responses were removed. The number of trials per condition was as follows (mean ± standard deviation): 67.9 ± 20 at rest, 53.5 ± 16.4 for uncertain go (early TMS), 53.0 ± 19.1 for certain go, 39.1 ± 13.4 for successful stop (late TMS), 46.4 ± 16.6 for uncertain go (late TMS), and 9.4 ± 6.9 for failed stop (late TMS) trials. MEP amplitude for each trial was calculated as the peak-to-peak difference within the 5 to 50 ms time window post-TMS. Resulting MEP amplitudes from valid trials were subsequently used for group-level analysis.

### Statistical analysis

Statistical analysis for TEP peaks was performed in MATLAB 2021b platform (MathWorks®, Inc., Massachusetts, USA) and R (v4.3.3). A paired-T test was applied to compare the TEP at rest for both targets. The two-way repeated measures analysis of variance (rANOVA) was applied to investigate the main effects (TMS type (active and sham) and Trial type (successful stops and uncertain goes for late TMS, and uncertain and certain go trials for early TMS)) and their interaction for TEP peak amplitude and task, respectively. Post-hoc test was conducted to further confirm interaction effects (Bonferroni correction p < 0.05). To investigate differences between active and sham TMS across the scalp, cluster-based permutation t-tests were applied across time points and electrodes, within each time window of interest (cluster forming threshold p<0.05, alpha<0.05 two-tailed, 1000 randomizations). For the correlation analysis, Pearson correlation coefficients were computed between TEP amplitude and SSRT in a similar manner. The scalp distributions were visualized using topographical maps. The Scheirer-Ray-Hare test (non-parametric rANOVA) was applied to compare the VAS rating. Pearson correlations were performed to test for a relationship between TEP peak amplitude, MEP amplitude (on M1) and the baseline task performance. The correlation between TEP peak amplitude and the VAS rating was performed using Spearman correlation analysis. The significance of results was set as p < 0.05 without correction.

## Results

### Behavioral performance

One participant was excluded from further analyses due to a low successful stop rate in the baseline task (< 30%) resulting in a final sample of 24 participants (age = 25.71 ± 4.37 years old, 13 females, 97.5 ± 6.08% right-handed). The overview of task performance is presented in Table 1. Omissions and errors for go trials were infrequent (typically below 2%) and did not show main effects of target site (M1 or preSMA) or TMS type (active or sham). Nevertheless, omission rates on uncertain go trials were significantly higher during active TMS in M1 than preSMA. The stop success rate was significantly lower during active than sham TMS and especially low for active M1 stimulation, possibly due to an involuntary finger movement induced by the TMS pulse.

**Table 1.**
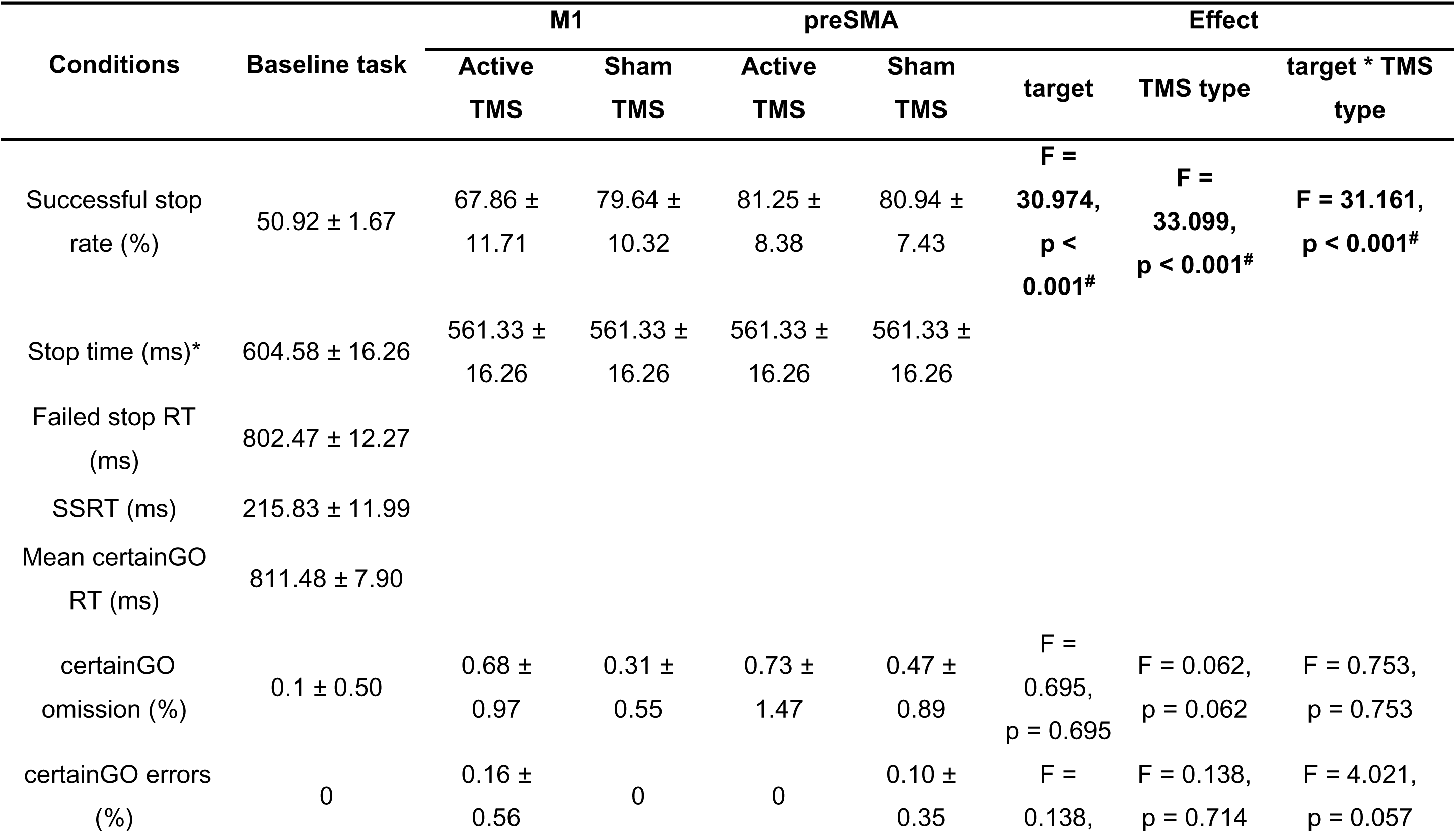
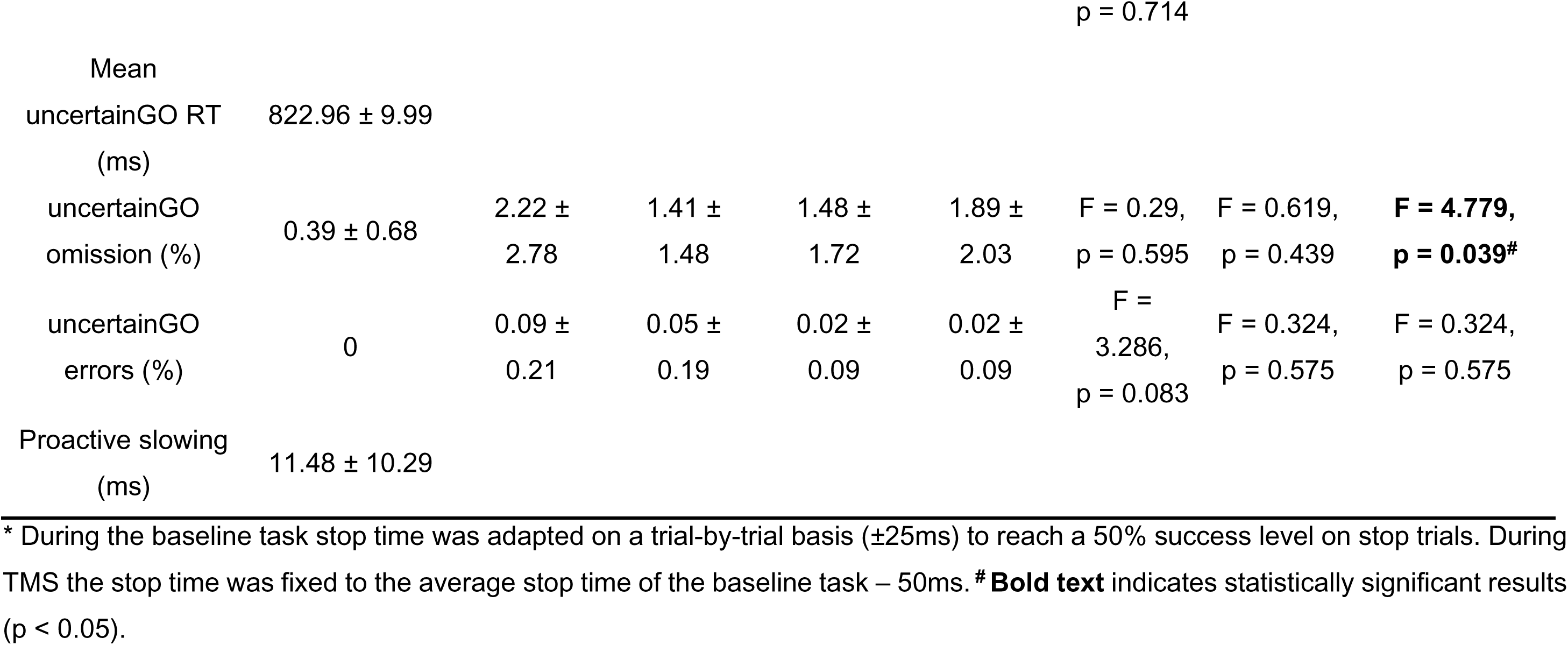
Task performance for different conditions

### VAS rating

All participants tolerated the stimulation well (49.88 ± 5.40 %MSO for M1 and 49.96 ± 5.31 %MSO for preSMA) as indicated by the low VAS scores with respect to pain (on average 2.15 for M1 and 1.45 for preSMA, Table S1). The sound of the TMS pulse was effectively masked in all conditions, with an average VAS score of 8.58 (between clearly and fully masked, Table S1). Importantly, there was no difference in the sound masking (H = 0.142, p = 0.706) or the intensity of the skin sensations (H = 0.905, p = 0.341) between the true and sham TMS blocks.

### TEP at rest

Active TMS on M1 increased N15, P30, P60 and P180 amplitude at rest. Active TMS on preSMA amplified N45 and N100 amplitude, in comparison to sham stimulation (Figure 2A, Table S2). Across participants, baseline SSRTs were associated with the difference in TEP amplitude between active and sham M1 stimulation. A larger N45 difference was related to faster SSRTs (r = 0.638, p = 0.002, Figure 2B). These relationships were mostly localized to the stimulation site (Figure 2C).

**Figure 2.**
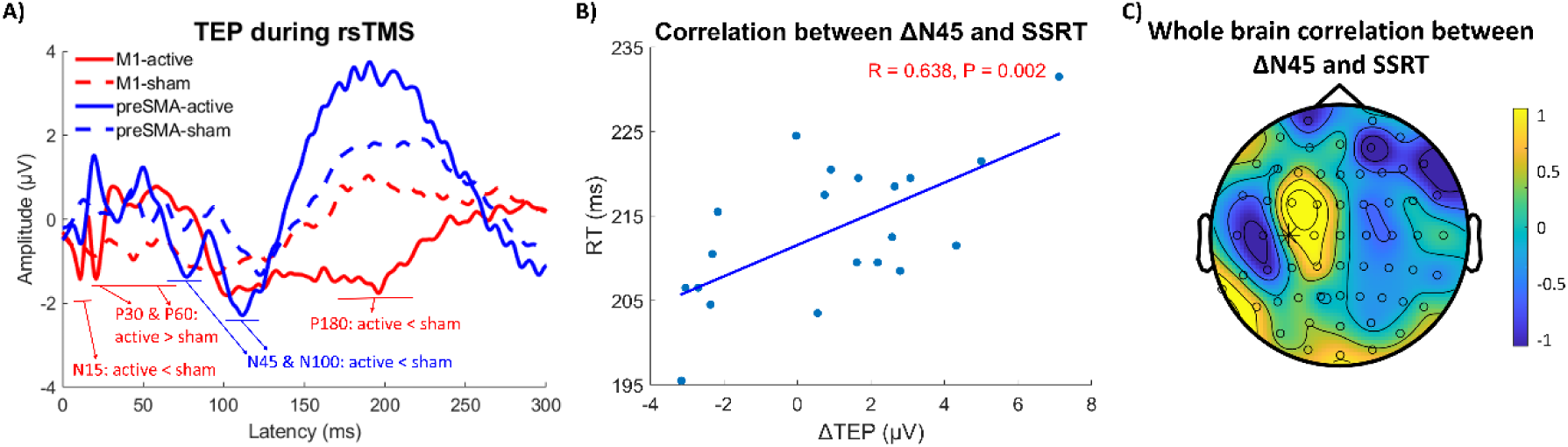
TEP during rsTMS (N = 22). **(A)** TEP time course and the peak amplitude comparison between active (solid line) and sham (dashed line) stimulation of channel C3 (i.e. close to site of M1 stimulation, red) and channel FC2 (i.e. close to site of preSMA stimulation, blue) at rest (p < 0.05). **(B)** Correlation between ΔN45 amplitude (active - sham) and baseline SSRT on M1 (C3) and for the **(C)** across the scalp. The yellow and blue represent positive and negative correlations in the topological plot. The star marks channel C3 for M1.

To further analyze the topographic distribution across the scalp, we conducted cluster-based permutation analysis between active and sham TMS conditions. There was no difference between active and sham conditions (Figure 3-4, top row).

**Figure 3.**
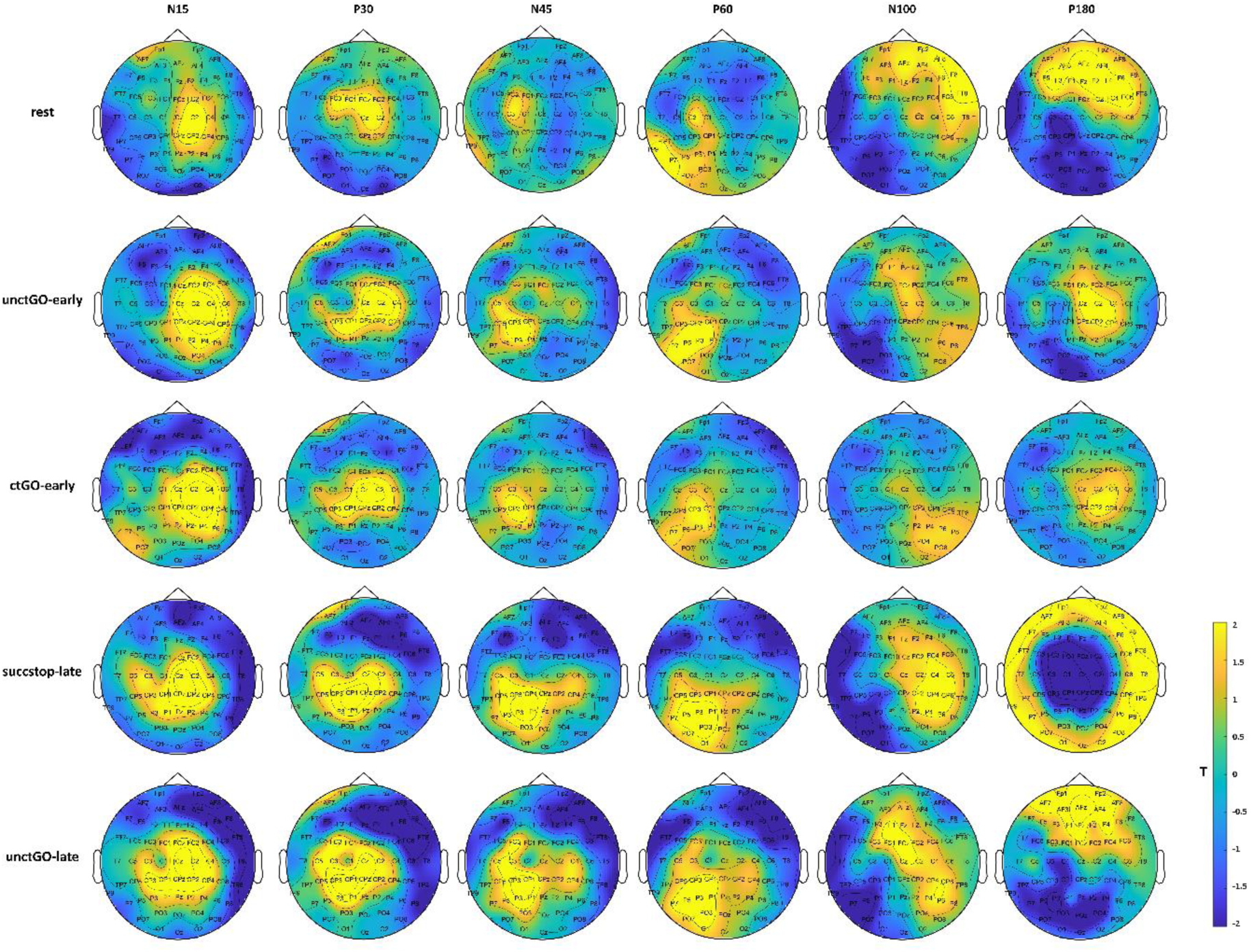
Results of cluster-based permutation analysis on M1 comparing active and sham stimulation. Trial type abbreviations: unctGO = uncertain go, ctGO = certain go, sucstop = successful stop.

**Figure 4.**
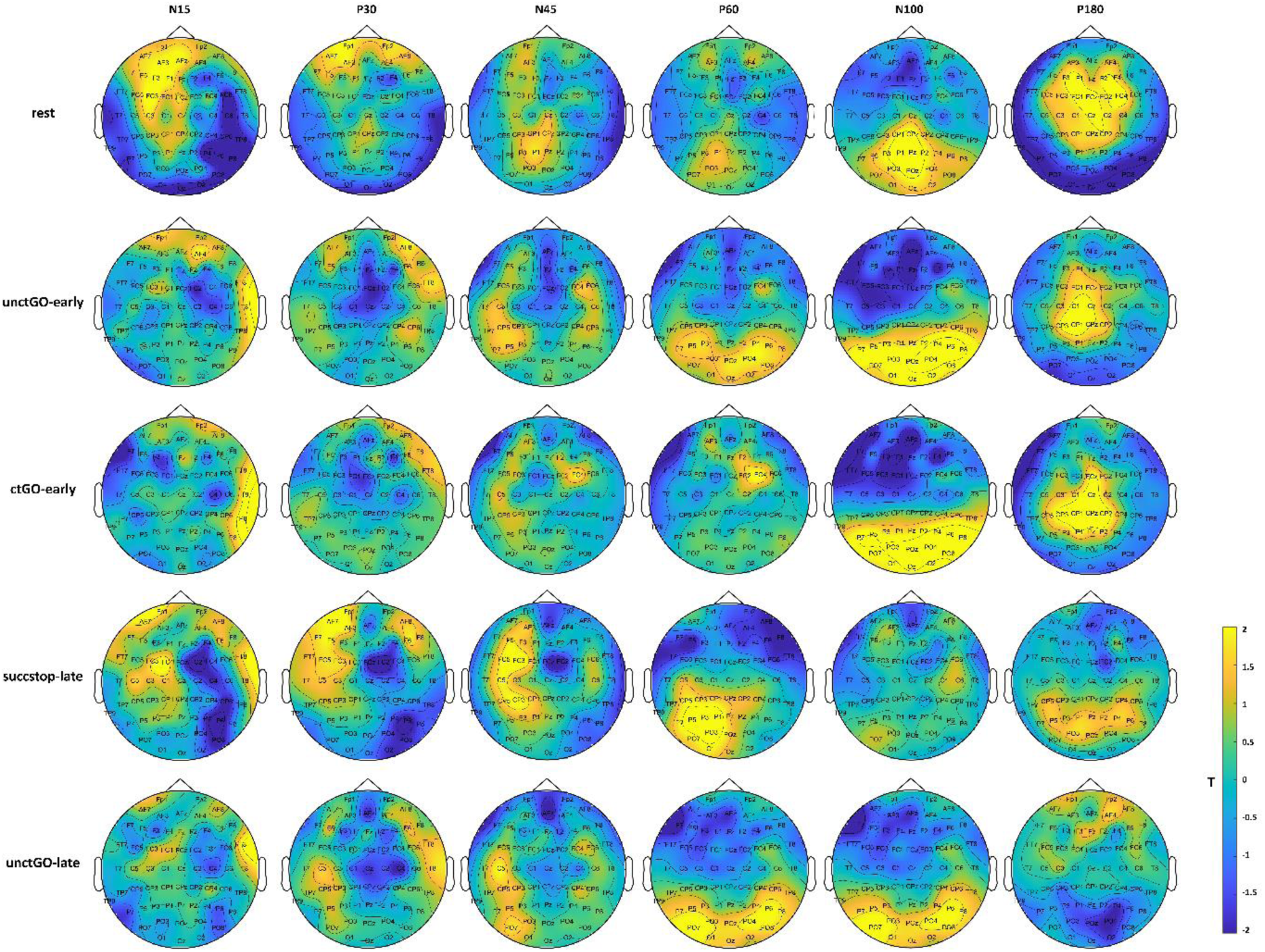
Results of cluster-based permutation analysis on preSMA. Trial type abbreviations: unctGO = uncertain go, ctGO = certain go, sucstop = successful stop.

### TEP during task performance

#### Early time point

Two participants were excluded from the early active TMS condition (i.e. 300ms after trial onset) on preSMA due to a strong baseline shift evoked by the TMS pulse, likely caused by the interaction between ongoing event-related activity in response to the start of the trial and the TMS evoked activity (Nikulin et al., 2007). Early M1 stimulation resulted in higher N15, and P60 amplitudes compared to sham stimulation (Figure 5A, Table 2). Similarly, early TMS on preSMA led to greater N15, P60 and higher P180 amplitudes under active stimulation compared to sham (Figure 5B). No main effects of Trial type were found for all TEP peaks. Only M1 P30 showed a significant TMS type and Trial type interaction. Certain go trials displayed a larger P30 amplitude during active stimulation compared to sham stimulation than uncertain go trials.

**Figure 5.**
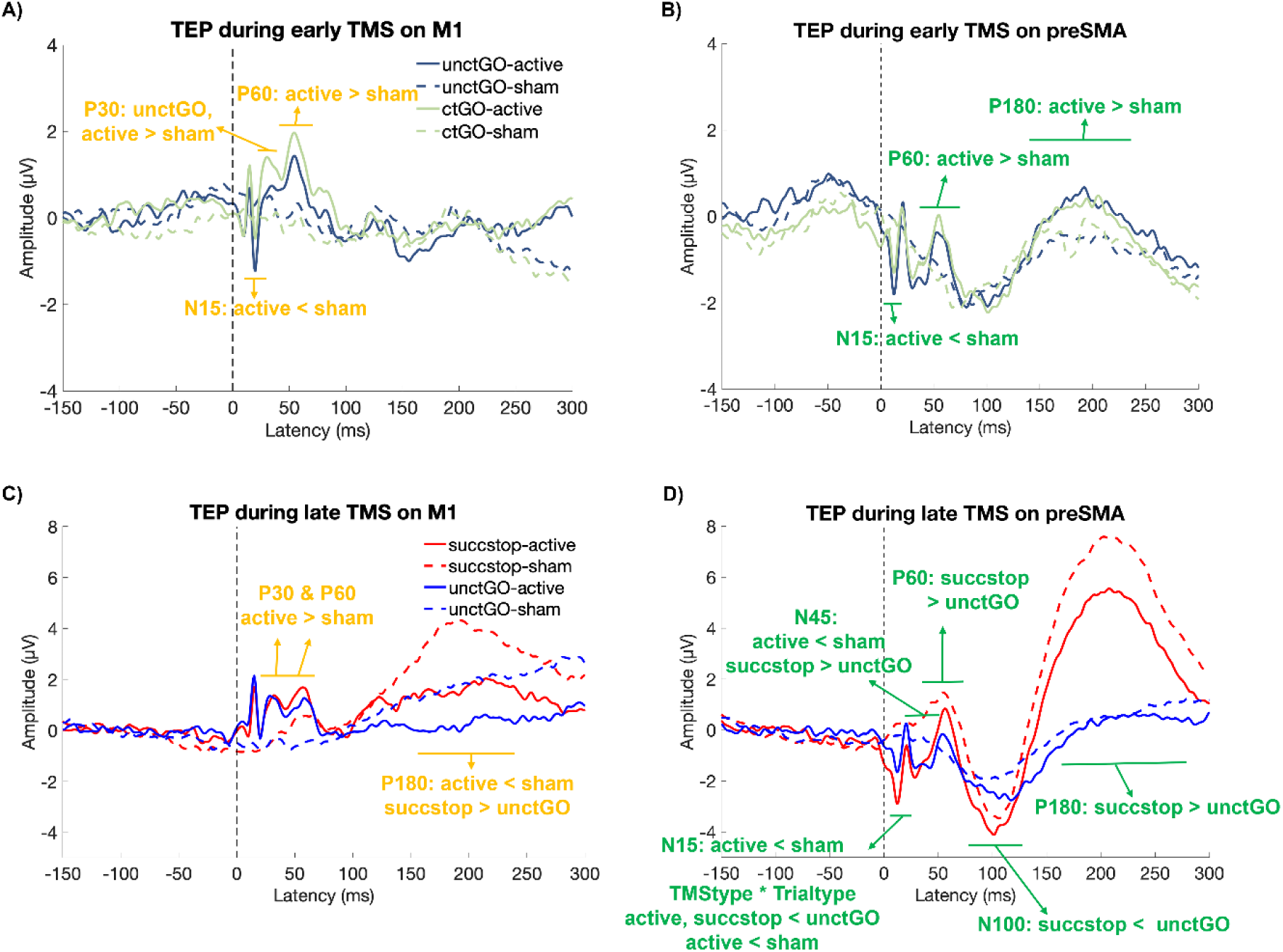
TEP during task performance (N = 24). The time course for early TMS on **(A)** M1 and **(B)** preSMA was plotted separately. Significant effects for rANOVA were highlighted in yellow for M1 and in green text for preSMA, consistent across all panels. The displays of the time course on **(C)** M1 and **(D)** preSMA under late TMS were shown in the lower panels.

**Table 2.**
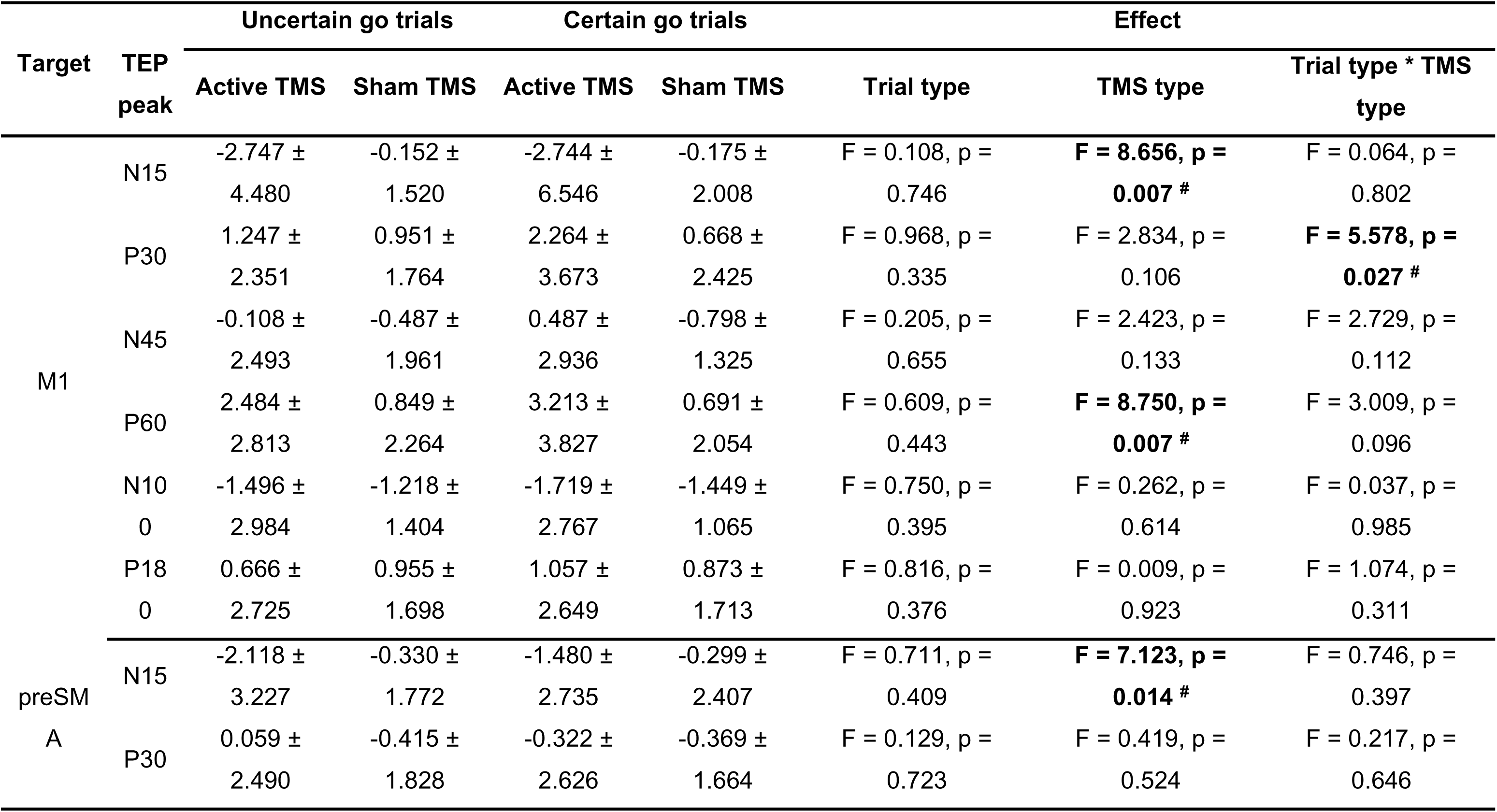
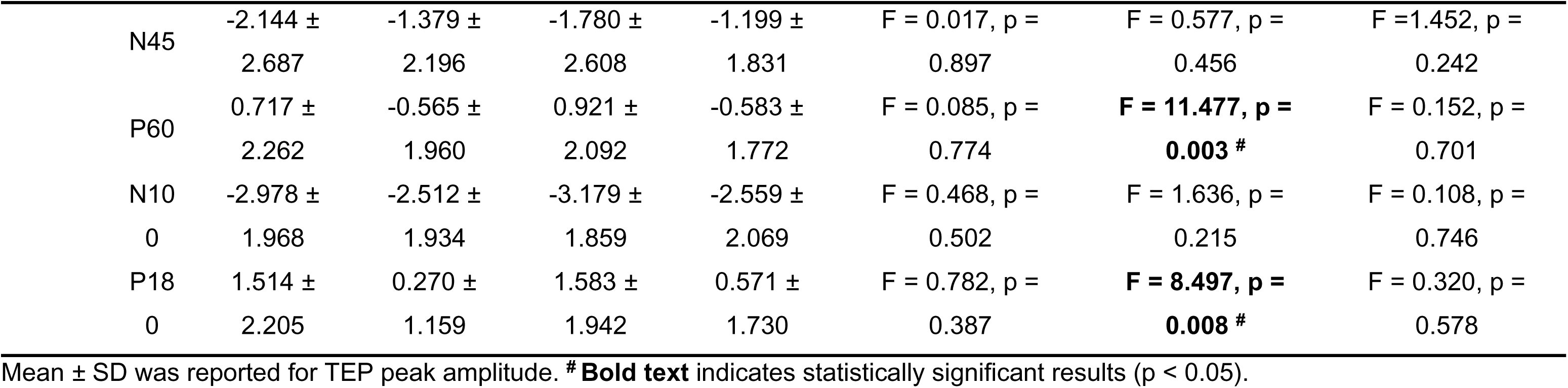
TEP peak amplitude for early TMS during task performance

However, this difference did not survive correction for multiple comparisons in post-hoc analyses.

#### Late time point

TMS targeting M1 showed a main effect of TMS type. Specifically, active late TMS on M1 increased P30 and P60 and lower P180 amplitude compared to sham (Figure 5C, Table 3). In addition, the P180 amplitude was significantly higher in successful stop trials compared to go trials. However, no interaction between Trial type and TMS type was found.

**Table 3.**
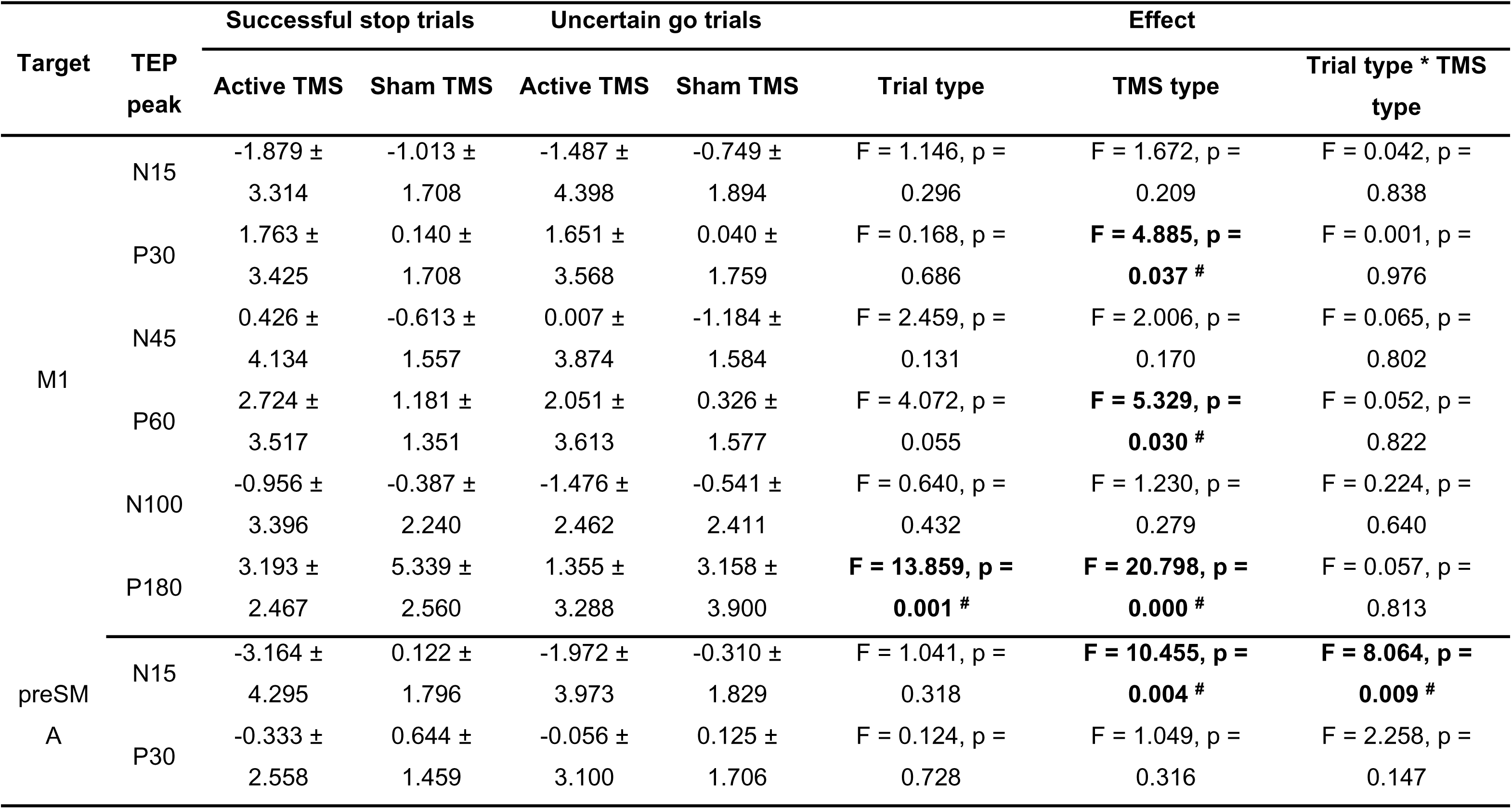
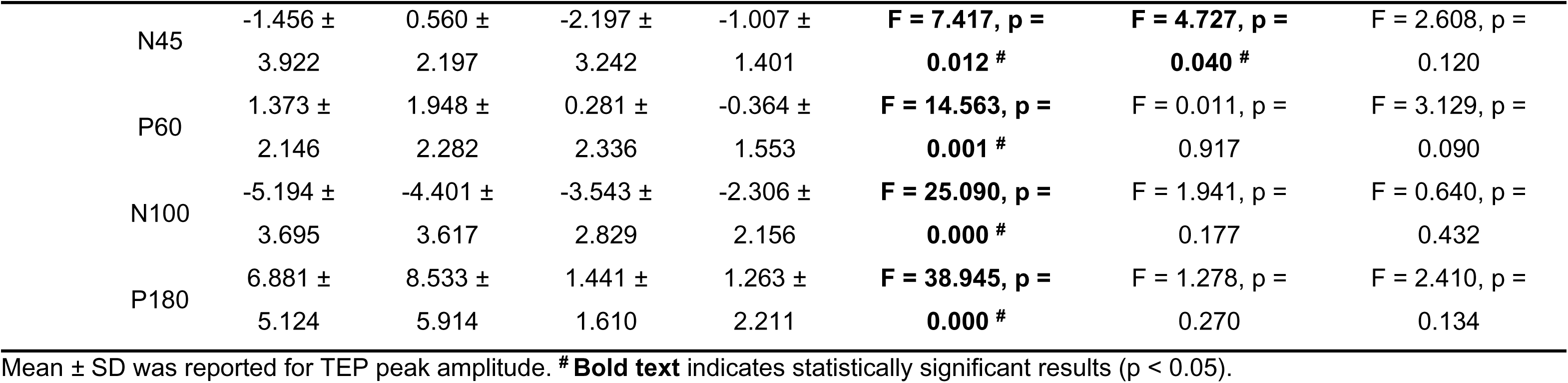
TEP peak amplitude for late TMS during task performance.

| Target | TEP peak | Successful stop trials |  | Uncertain go trials |  | Effect |  |  |
| --- | --- | --- | --- | --- | --- | --- | --- | --- |
|  |  | Active TMS | Sham TMS | Active TMS | Sham TMS | Trial type | TMS type | Trial type * TMS type |
| M1 | N15 | -1.879 ± | -1.013 ± | -1.487 ± | -0.749 ± | F = 1.146, p = | F = 1.672, p = | F = 0.042, p = |
|  |  | 3.314 | 1.708 | 4.398 | 1.894 | 0.296 | 0.209 | 0.838 |
|  | P30 | 1.763 ± | 0.140 ± | 1.651 ± | 0.040 ± | F = 0.168, p = | **F = 4.885, p =** | F = 0.001, p = |
|  |  | 3.425 | 1.708 | 3.568 | 1.759 | 0.686 | **0.037 #** | 0.976 |
|  | N45 | 0.426 ± | -0.613 ± | 0.007 ± | -1.184 ± | F = 2.459, p = | F = 2.006, p = | F = 0.065, p = |
|  |  | 4.134 | 1.557 | 3.874 | 1.584 | 0.131 | 0.170 | 0.802 |
|  | P60 | 2.724 ± | 1.181 ± | 2.051 ± | 0.326 ± | F = 4.072, p = | **F = 5.329, p =** | F = 0.052, p = |
|  |  | 3.517 | 1.351 | 3.613 | 1.577 | 0.055 | **0.030 #** | 0.822 |
|  | N100 | -0.956 ± | -0.387 ± | -1.476 ± | -0.541 ± | F = 0.640, p = | F = 1.230, p = | F = 0.224, p = |
|  |  | 3.396 | 2.240 | 2.462 | 2.411 | 0.432 | 0.279 | 0.640 |
|  | P180 | 3.193 ± | 5.339 ± | 1.355 ± | 3.158 ± | **F = 13.859, p =** | **F = 20.798, p =** | F = 0.057, p = |
|  |  | 2.467 | 2.560 | 3.288 | 3.900 | **0.001 #** | **0.000 #** | 0.813 |
| preSM | N15 | -3.164 ± | 0.122 ± | -1.972 ± | -0.310 ± | F = 1.041, p = | **F = 10.455, p =** | **F = 8.064, p =** |
|  |  | 4.295 | 1.796 | 3.973 | 1.829 | 0.318 | **0.004 #** | **0.009 #** |
|  | P30 | -0.333 ± | 0.644 ± | -0.056 ± | 0.125 ± | F = 0.124, p = | F = 1.049, p = | F = 2.258, p = |
|  |  | 2.558 | 1.459 | 3.100 | 1.706 | 0.728 | 0.316 | 0.147 |

|  |  |  |  |  |  |  |  |
| --- | --- | --- | --- | --- | --- | --- | --- |
| N45 | -1.456 ± | 0.560 ± | -2.197 ± | -1.007 ± | **F = 7.417, p =** | **F = 4.727, p =** | F = 2.608, p = |
|  | 3.922 | 2.197 | 3.242 | 1.401 | **0.012 #** | **0.040 #** | 0.120 |
| P60 | 1.373 ± | 1.948 ± | 0.281 ± | -0.364 ± | **F = 14.563, p =** | F = 0.011, p = | F = 3.129, p = |
|  | 2.146 | 2.282 | 2.336 | 1.553 | **0.001 #** | 0.917 | 0.090 |
| N100 | -5.194 ± | -4.401 ± | -3.543 ± | -2.306 ± | **F = 25.090, p =** | F = 1.941, p = | F = 0.640, p = |
|  | 3.695 | 3.617 | 2.829 | 2.156 | **0.000 #** | 0.177 | 0.432 |
| P180 | 6.881 ± | 8.533 ± | 1.441 ± | 1.263 ± | **F = 38.945, p =** | F = 1.278, p = | F = 2.410, p = |
|  | 5.124 | 5.914 | 1.610 | 2.211 | **0.000 #** | 0.270 | 0.134 |

Mean ± SD was reported for TEP peak amplitude. # **Bold text** indicates statistically significant results (p < 0.05).

In contrast, late TMS on preSMA primarily elicited the main effect of Trial type, with larger P60, N100 and P180 amplitudes and smaller N45 observed in successful stop trials than uncertain go trials (Figure 5D). The N15 amplitude exhibited a main effect of TMS type, with greater amplitude during active stimulation. Notably, an interaction between Trial type and TMS type was observed for N15. Post hoc analysis indicated a Trial type difference solely under active stimulation, with greater N15 amplitude in successful stop than uncertain go trials (p = 0.05). Moreover, within successful stop trials, the N15 amplitude was significantly higher under active TMS compared to sham stimulation (p < 0.001).

#### Topographic distribution

To further explore the topographic distribution of the potentials across the scalp, we performed a cluster-based permutation analysis comparing active and sham TMS conditions for each trial type around the time points that the TEP peaks appeared (Figures 3-4). In line with previous findings, the analysis revealed that early TEPs were more localized to the stimulation target, whereas later TEPs showed a more widespread distribution across the cortex (Biabani et al., 2019; Rogasch et al., 2020). Yet, cluster-based t-statistics did not reveal significant TEP amplitude differences outside of the stimulation area.

### ERP for no-stimulation trials during task

Prominent P2, N2 and P3 peaks were observed following the stop-signal for both M1 and preSMA (Figure 6, Table S3). Notably, there is a high degree of temporal overlap between the stop-signal related P2 and TEP P60, between the N2 and TEP N100, and between the P3 and the TEP P180 (Figure 6). Despite this overlap, no significant correlation was found between the corresponding TEP and ERP amplitudes. The rsTEP showed an intriguingly similar pattern to the taskTEP and ERP on preSMA. This suggested that the preSMA may have a stable baseline excitability or network state that does not dramatically change regardless of the task state.

**Figure 6.**
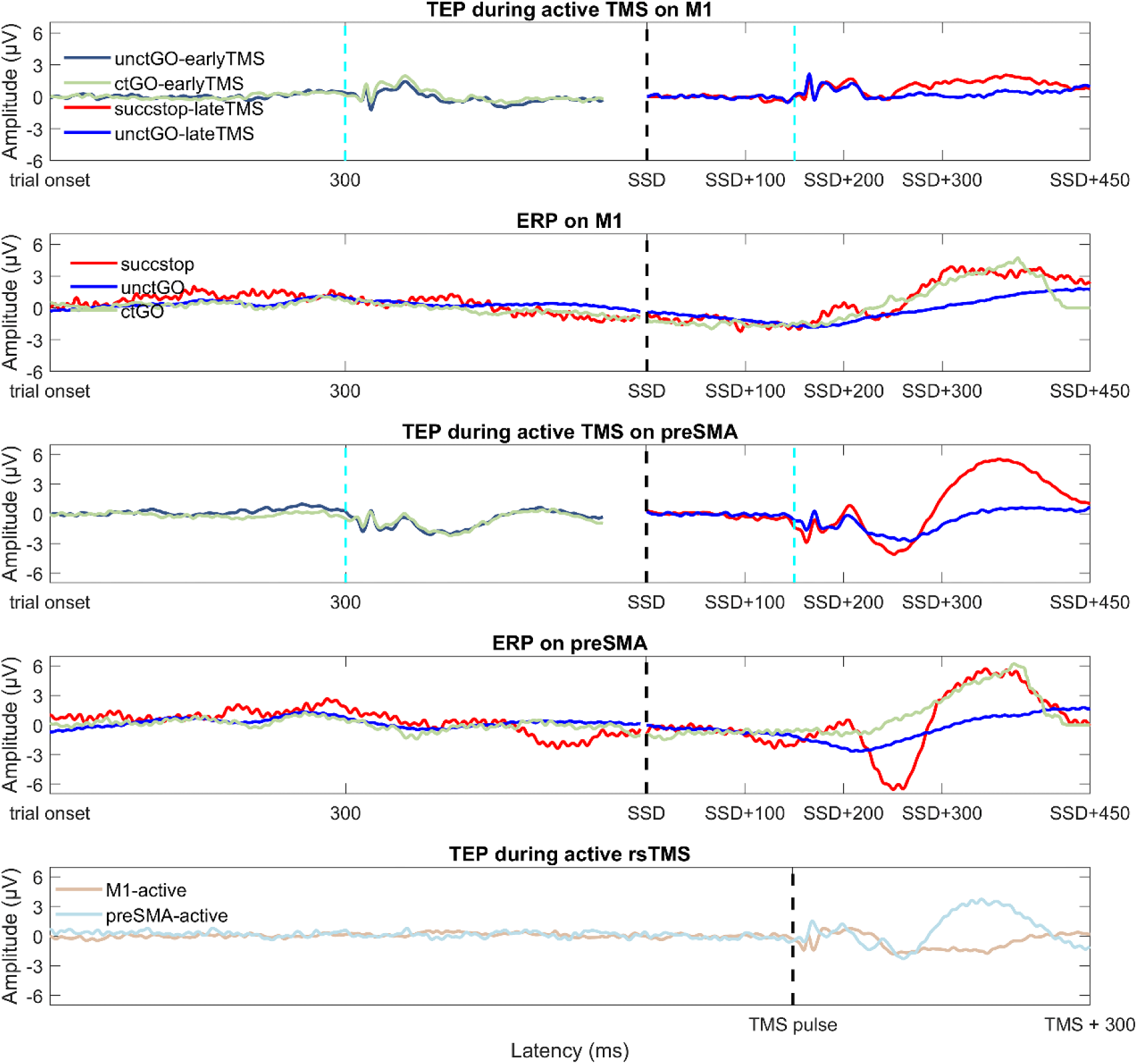
ERP and TEP time course in the time domain. The time courses for TEP and ERP under different conditions were aligned to start at the trial onset. During task performance, early TMS was applied 300ms after trial onset, while late TMS was delivered 150ms after individual SSD. The TMS pulse was indicated by cyan dotted lines, and the group average SSD for stop trials was represented by black dotted lines. The TEP components P60, N100, and P180 corresponded to ERP components P2, N2, and P3, respectively. In terms of the TEP at rest, the TMS pulse was aligned with late TMS. The rsTEP in preSMA mirrored the pattern observed during task performance. SSD = stop-signal delay.

### MEP during TMS on M1

The MEP results indicated that MEP amplitudes showed a main effect (F = 32.078, p < 0.001) for trial types. MEP at rest were lower compared to task conditions (Figure 7). Among the task conditions, failed stop and uncertain go trials (during late TMS) elicited the largest MEP amplitudes, significantly exceeding those observed in other task conditions. However, we did not find any correlation between MEP and TEP, nor between MEP and inhibitory performance.

**Figure 7.**
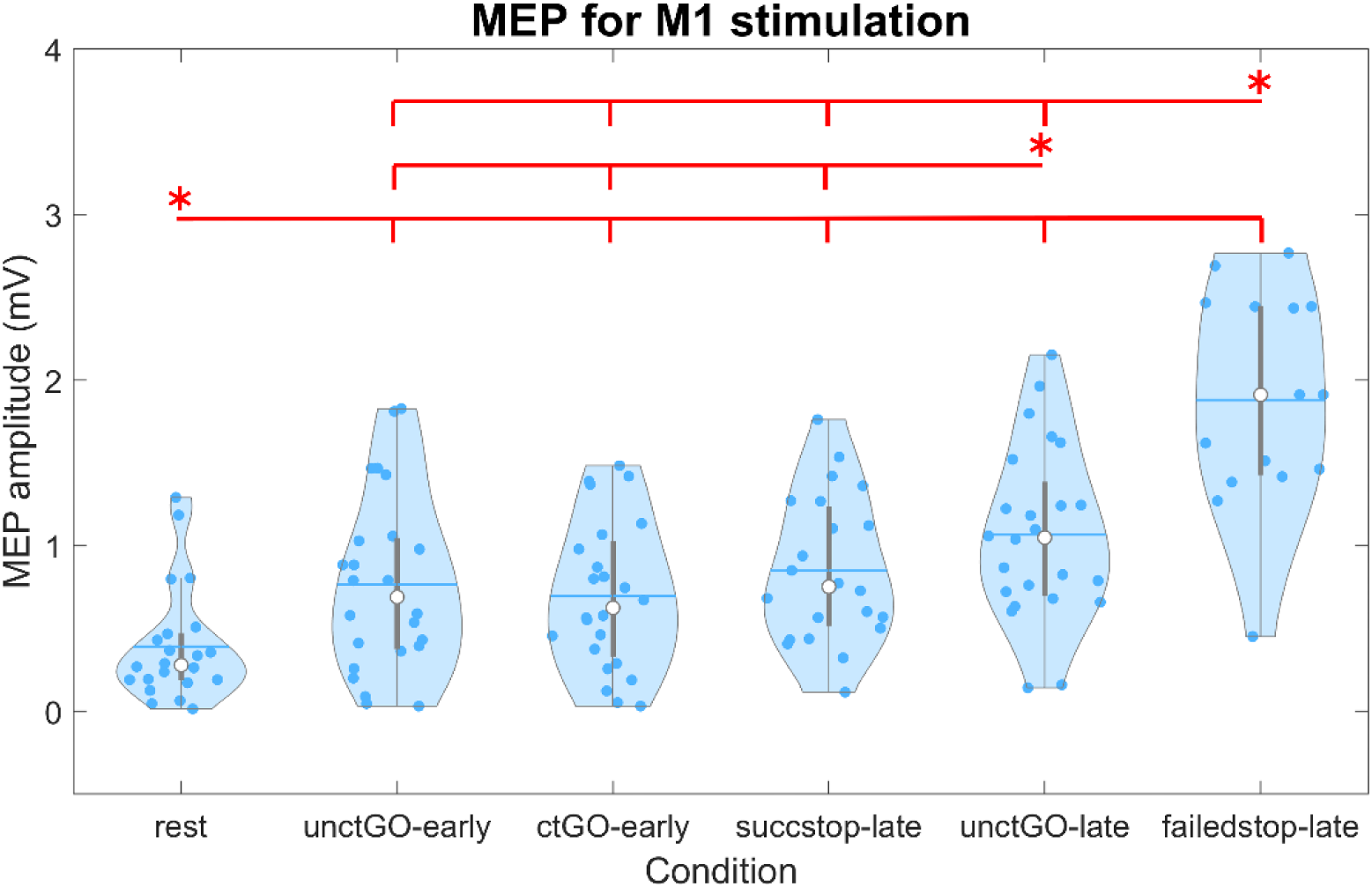
MEP amplitudes across conditions during M1 stimulation. The violin plot displays the distribution of MEP amplitudes, with black and red lines indicating the mean and median at the group level, respectively. Grey scatter points represent individual data values for each condition. Red horizontal lines denote statistically significant differences between conditions, as determined by pairwise comparisons. Asterisks (*) mark conditions that are significantly different from all other conditions connected by the corresponding red line (p < 0.05). Trial type abbreviations: unctGO = uncertain go, ctGO = certain go, sucstop = successful stop, failedstop = failed stop.

These findings demonstrated that motor excitability was modulated by task demands. The higher MEPs observed in uncertain go and failed stop trials may be attributed to a combination of cortical readiness in response to uncertainty, resulting in increased cognitive and motor effort (Alhussein & Smith, 2021).

## Discussion

This study combined TMS-EEG to probe GABA- and glutamatergic signaling during reactive and proactive motor inhibition. Importantly, compared to ppTMS, spTMS does not require a substantial delay between pulses (e.g. LICI), making it possible to study cortical inhibition as the reactive inhibition process unfolds. In addition, TMS-EEG allows to investigate receptor activity across the cortex, not being restricted to M1. Here we also targeted preSMA, providing a more complete understanding of GABAergic and glutamatergic signaling in the wider motor inhibition network during reactive and proactive inhibition.

### N45 amplitude at rest predicts reactive inhibition performance

The relationship between M1 N45 peak amplitude at rest and SSRT suggests that GABAa supports reactive inhibition. This finding is in line with previous studies demonstrating that individuals with higher rest SICI are more efficient at inhibiting prepotent actions (Cardellicchio et al., 2021; Chowdhury et al., 2018; Loomes et al., 2023; Tran et al., 2020). Opposed to TMS-EMG, it is possible to look beyond M1 with TMS-EEG. As typically observed N45 amplitudes were greatest in centro-frontal electrodes (Figure 3) (Gordon et al., 2021). Although N45 amplitudes did not significantly differ between active and sham TMS on C3, it was specifically the N45 recorded over C3 that correlated with behavioral performance (SSRT). The whole-brain correlation plot further confirmed that the relation between N45 and SSRT was confined to sensory-motor regions.

### Early TEP peak N15 is larger during successful stopping

To assess changes in TEP amplitude during successful reactive inhibition TMS was applied 150ms after the stop-signal. During this time window, inhibition is implemented and starts to become evident as reduced MEP amplitude in the effector agonist muscle (Coxon et al., 2006; Jana et al., 2020; MacDonald et al., 2014). N15 peak amplitude was significantly larger during successful stopping when active TMS was applied over preSMA. This early TEP component is typically localized over and near the target site (Bonato et al., 2006; Darmani & Ziemann, 2019; Esser et al., 2006; Komssi et al., 2004; Maki & Ilmoniemi, 2010; ter Braack et al., 2015) and is believed to reflect excitation (Maki & Ilmoniemi, 2010; ter Braack et al., 2015). The preSMA plays a crucial role in reactive stopping by rapidly sending excitatory signals to the STN via the hyperdirect pathway, ultimately, leading to the suppression of motor output (Jahanshahi et al., 2015; Wolpe et al., 2022). We hypothesize that stronger early excitation in the preSMA enhances the initiation of inhibitory processes mediated by cortico-subcortical interaction.

In contrast, no interaction was found for the N45 or N100 peak amplitude in either preSMA or M1, suggesting that cortical inhibitory processes did not significantly increase during reactive inhibition. This finding is particularly surprising for the N45. M1 N45 amplitude at rest correlated with SSRT, and previous research has shown that short-interval intracortical inhibition (SICI) increases during successful stopping at the same timepoint (Coxon et al., 2006; Hermans et al., 2019). One possible explanation is that task-TMS interactions at these later timepoints are obscured by temporal overlap in the TEP and event-related potentials (ERPs) generated in response to the stop-signal and/or peripherally evoked potentials (PEPs) from auditory and somatosensory processing of the TMS pulse (see following section for a more in-depth discussion).

It is also important to note that the relationship between TEPs and GABAergic/glutamatergic activity is largely inferred from pharmacological studies (Belardinelli et al., 2021; Darmani et al., 2016; Premoli et al., 2014b; Rogasch et al., 2020) and does not directly establish causality. For instance, studies using double-pulse TMS-EEG have demonstrated that the pharmacological effects of GABAergic modulators differ between double-pulse MEPs and TEPs (Premoli et al., 2018; Premoli et al., 2014b), underscoring the challenges in using these indirect measures to investigate the activity of specific neurotransmitter receptors. In addition, it is also still largely unknown to what extent the pharmacological characterization of TEPs in M1 applies to TEPs elicited in other areas (Salavati et al., 2018; Song et al., 2025).

### Disentangling TMS evoked potential from other evoked responses

Despite the limited amount of trials without TMS stimulation (20 stop trials and 100 go trials), we were able to identify clear P2, N2 and P3 ERP components as typically observed in stop compared to go trials (Huster et al., 2020; Kok et al., 2004). Comparing the ERP with the TEP in the preSMA, and to a lesser extent also on M1, reveals a clear overlap in peak timings from approximately 50ms after the TMS pulse, i.e., 200ms after the stop signal was presented (Figure 6). There also seem to be only task induced effect in the TEP (stop > go) from this moment onwards (∼50ms after the TMS pulse) and no differences between sham and active TMS (Figure 6). This suggests that the evoked response to the stop-signal was so strong that any small TMS effect became unobservable. It also raises the question whether the TEP during successful stopping and the ERP might share generating mechanisms, such as thalamocortical dynamics (Diesburg et al., 2023), making it difficult to fully disentangle TMS evoked potentials from ongoing event-related responses during a task.

Furthermore, the multisensory inputs from the TMS pulse, including auditory and somatosensory stimuli, generate evoked responses starting around 100 ms after the pulse (Ilmoniemi & Kicic, 2010). In this study, we attempted to minimize the contamination from these peripherally evoked potentials (PEPs) by using a sham procedure that delivered the same multisensory stimuli as active TMS. Participants’ subjective ratings confirmed that the perceived intensity of the TMS pulse and scalp sensations were comparable between sham and active TMS conditions. However, differences in the N100 and P180 TEP peak amplitudes at rest, particularly in the preSMA, suggest that this approach may have been insufficient to fully eliminate PEPs, and that later TEP components may still be influenced by them. Song et al. (2024) observed comparable PEPs in the SMA for both sham and active TMS using a procedure very similar to ours (Song et al., 2024). The main distinction was the intensity of electrical stimulation—24 mA in their study compared to an average of 18 mA in ours. It is possible that higher stimulation intensity is required to fully saturate the EEG response to sensory inputs. Nevertheless, since the sensory stimulation is identical across task conditions during active TMS, any differences in TEPs between trial types can be attributed to direct cortical stimulation, as long as the experimental manipulation does not evoke differential ERPs (e.g. stop vs go) or alter sensory input processing.

Finally, the contrast in TEPs between stop and uncertain go trials in M1 could be influenced by differences in the strength of re-afferent somatosensory evoked potentials. TMS pulses applied at intensities above the resting motor threshold can trigger a motor response, which is followed by re-afferent feedback from the muscle to the motor cortex. Our MEP analysis demonstrated that MEP amplitudes are significantly lower in successful stop trials compared to uncertain go trials, suggesting that less re-afferent feedback reaches the motor cortex during successful stopping. Previous studies have shown that TEP amplitudes diverge when comparing trials with and without MEPs following a TMS pulse at 100% resting motor threshold, with differences emerging as early as ∼45 ms post-stimulation (Biabani et al., 2021; Fecchio et al., 2017; Petrichella et al., 2017). In light of our findings, the lack of a TEP peak amplitude difference between successful stop and uncertain go trials may be explained by an inflated TEP amplitude in the go trials due to stronger re-afferent feedback. This increased feedback likely masks the true differences in cortical excitability between conditions. To mitigate this effect, future studies should consider using subthreshold stimulation intensities, calibrated in the active muscle, to minimize re-afferent contributions and allow for a clearer assessment of TEPs related to cortical processes.

Due to all the above, the results for TEPs at the late stimulation time point should be considered inconclusive from ∼45ms onwards, as the combination of strong task-related evoked responses, remaining peripheral evoked responses and re-afferent feedback likely obscure the true TMS evoked dynamics during motor inhibition.

### Reduced motor cortex facilitation during proactive inhibition

When there is a possible need for motor inhibition, participants tend to slow down in order to increase the likelihood of being able to inhibit their movement. In the present study, the response times of go trials in which a stop-signal could appear compared to certain go trials showed the expected proactive slowing (Vink et al., 2015). Previous TMS studies utilizing a paired-pulse paradigm over M1 found higher LICI in go trials where stopping might be required (Hermans et al., 2019). Here, we observed significant differences in peak amplitude between active and sham TMS on M1 and preSMA, but there was no difference between certain and uncertain go trials or an interaction. Only the M1 P30 peak amplitude showed a significant interaction between TMS type and trial type, with higher P30 amplitudes for certain than uncertain go trials under active TMS. This pattern may reflect reduced motor facilitation during proactive inhibition.

In rest LICI relates to the slope of the N100 (Opie et al., 2017; Rogasch et al., 2013), therefore one would expect higher N100 amplitudes during uncertain versus certain go trials. This null result cannot be explained by event-related potential clouding the TEP (Figure 6). Therefore, it raises the question whether LICI and N100 reflect the same inhibitory activity in this task context. Given the overlap between the N100 and the auditory PEP, previous findings linking the N100 to GABAergic activity have been called into question. Recently, Gordon et al. (2023b) demonstrated that GABAa agonists increase N100 amplitude during both active and sham TMS, suggesting that the modulation observed in the N100 may be influenced by PEPs rather than reflecting direct GABAergic activity.

### Limitations and future directions

Despite applying white noise and CES to mask auditory and somatosensory inputs, the sham procedure may not have fully replicated the sensory experience of active TMS. Even though VAS scores did not indicate less effective masking, differences in N100 and P180 peaks between active and sham rest TEPs were observed, particularly in the preSMA, which implies that the TEPs elicited by sham and active TMS were not equally matched. Perhaps a higher CES amplitude is required to saturate the engaged cranial nerves and/or a more personalized titration of the stimulation intensity. These findings highlight the ongoing challenge of achieving equivalence between active and sham conditions. Improved sham protocols, such as advanced coil designs, enhanced sensory-masking techniques and target specific calibration, are essential to address this limitation.

The assumption underlying the approach to provide CES during both sham and active TMS is that the resulting TEPs and PEPs are linearly superimposed. It is however not inconceivable that the high-intensity peripheral stimulation interacts with the brain response to TMS. Recent evidence demonstrates that, at least at the macroscopic EEG level, there is no evidence for such an interaction (Gordon et al., 2023a).

In terms of timing, administering TMS pulses 150ms after the stop signal on both M1 and preSMA might have been a bit late for preSMA. Jana et al. (2020) reported that stop command from preSMA might be transferred to motor cortex ∼120ms after stop signal, suggesting that an even stronger difference in TEP peak amplitudes between go and stop trials might be observed when stimulating slightly earlier (Jana et al., 2020). Finally, the stop signals on TMS trials were presented 50ms earlier than individual stop signal delays during baseline task performance. This adjustment was made to ensure more successful stop trials for TEP analysis and to avoid stimulating after movement onset in go trials. However, by making the task easier, participants might have engaged in a less reactive stopping process than they would with stop signal delays that matched their SSRT, potentially influencing the results.

## Conclusion

The present study provides novel insights into the neural mechanisms of motor inhibition by combining TMS with EEG. Unlike conventional TMS-EMG measures that solely focus on M1, TMS-EEG enables investigation of GABA- and glutamatergic receptor involvement across the cortex. By stimulating preSMA, a key node within the motor inhibition network, we observed increased N15 amplitudes during both reactive and proactive inhibition. This likely reflects fast excitatory signals from preSMA towards the basal ganglia to initiate or implement motor inhibition. Additionally, P30 peak amplitudes were higher for certain than uncertain go trials in M1, potentially indicating a reduction in motor facilitation during proactive inhibition.

## Data and Code Availability

The data that support the findings of this study are available on request from the corresponding author, I.L.

## Author contribution

T.Z.: Data Curation, Formal analysis, Investigation, Methodology, Software, Validation, Visualization, Writing - original draft, Writing - review & editing. C.C.: Investigation, Methodology, Software, Writing - review & editing. J.C.: Conceptualization, Methodology, Writing - Review & Editing. A.T.S.: Funding acquisition, Methodology, Supervision, Writing - review & editing. I.L.: Conceptualization, Formal analysis, Funding acquisition, Investigation, Methodology, Software, Supervision, Validation, Writing - review & editing.

