## Supplementary_materials for "Using TMS-EEG to study the intricate interplay between GABAergic inhibition and glutamatergic excitation during reactive and proactive motor inhibition"

*Corresponding author

### Masking procedure for control condition

#### Sound masking procedure

The waveform of the TMS click was digitized and processed to produce a continuous audio signal that captured the specific time-varying frequencies by using a Matlab script (2020b) with NI-DAQ hardware. The volume of the white noise gradually increased up to maximally 2.5 Vrms (mean = 2.16 ± 0.41 Vrms) until the 'click' sound from TMS could no longer be perceived or until they had reached their threshold for comfort (always below 100 dB). The determination of sound intensity was applied using PsychoPy (v2022.2.5). The sound intensity given to both ears was ensured to be the same through the digital setting and participants' feedback.

#### Sensory masking procedure

Two bipolar electrodes were placed near the TMS target site (8mm in diameter, the cross points of the surrounding electrodes from C3 were used for M1, and from FC2 for preSMA – to deliver CES. The pulse consisted of a biphasic square wave pulse of 50µs duration and 200V compliance voltage. The intensity was set individually at 300% of the lowest intensity at which participants could perceive the stimulation (mean stimulation intensity = 18.3 ± 5.5mA) (Conde et al., 2019; Gordon et al., 2021; Lin et al., 2003; Torquati et al., 2002).

### Supplementary methods

#### VAS comparison

The sound of the TMS pulse was effectively masked in all conditions, with an average VAS score 8.58 (between clearly and fully masked, Table S1). Importantly, there was neither a main effect of target nor TMS type for sound masking and stimulation area (Table S1). However, participants reported lower sensation (skin sensation and painful sensation) on preSMA than M1. There was no interaction between target and TMS type for all questions (Table S1). When asking the participants explicitly for the difference between the stimulation blocks after the experiment, one participant guessed there was sham condition, and three out of twenty-five participants reported TMS intensity changes.

### ERP comparison between successful stop and go trials

During successful stopping, the ERP P2, N2 and P3 were greater than uncertain go trials on M1 and preSMA, except the N2 on M1. Interestingly, there was no significant correlation between the late TEP components and ERP (Figure S1, Table S2).

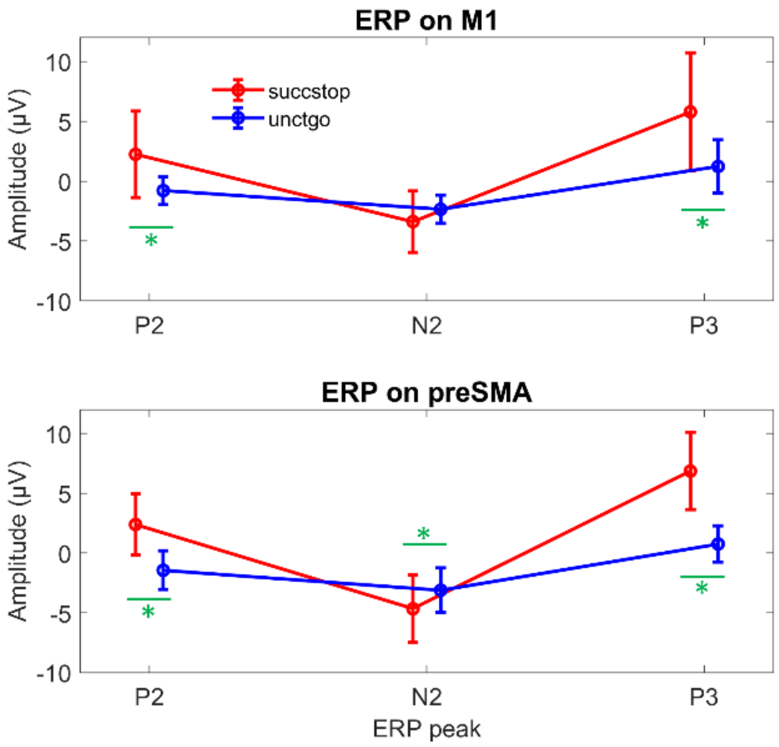


Figure S1. ERP peak amplitude during task performance. The green star and line represent the significant difference between successful stop and uncertain go trials ($p < 0.05$).

Table S1. Scheirer-Ray-Hare test for VAS

| items | M1 | | | preSMA | | | effect | | |
|---|---|---|---|---|---|---|---|---|---|
| | active TMS | sham TMS | active - sham | ative TMS | sham TMS | active - sham | target | TMS type | target * TMS type |
| Sound mask | 8.64 ± 1.78 | 8.52 ± 1.68 | 0.11 ± 0.92 | 8.53 ± 2.19 | 8.63 ± 1.89 | -0.09 ± 0.75 | H = 0.016, p = 0.900 | H = 0.142, p = 0.706 | H = 0.002, p = 0.965 |
| Skin sensation | 4.16 ± 1.6 | 4.2 ± 1.64 | -0.05 ± 0.95 | 3.08 ± 1.75 | 3.89 ± 1.84 | -0.81 ± 1.78 | **H = 4.288, p = 0.038**[#] | H = 0.905, p = 0.341 | H = 0.938, p = 0.333 |
| Stimulated area | 2.59 ± 0.97 | 2.95 ± 1.24 | -0.36 ± 0.71 | 2.57 ± 0.99 | 2.75 ± 1.13 | -0.18 ± 0.68 | H = 0.435, p = 0.510 | H = 0.753, p = 0.385 | H = 0.081, p = 0.776 |
| Painful sensation | 2.09 ± 1.76 | 2.2 ± 1.57 | -0.11 ± 0.79 | 1.41 ± 1.37 | 1.5 ± 1.59 | -0.09 ± 0.75 | **H = 5.286, p = 0.022**[#] | H = 0.150, p = 0.698 | H = 0.114, p = 0.736 |

Mean ± SD was reported for VAS scores. [#] **Bold text** indicates statistically significant results ($p < 0.05$)

Table S2. TEP at rest

| Target | TEP peak | Active TMS | Sham TMS | Active vs sham TMS |
|---|---|---|---|---|
| M1 | N15 | -3.95 ± 9.19 | -0.91 ± 2.53 | **T = -2.751, p = 0.012**[#] |
| | P30 | 1.93 ± 5.22 | 0.10 ± 1.83 | **T = 3.010, p = 0.007**[#] |
| | N45 | -0.59 ± 1.82 | -1.19 ± 1.02 | T = 1.300, p = 0.208 |
| | P60 | 2.50 ± 2.36 | 0.32 ± 0.80 | **T = 4.170, p = 0.000**[#] |
| | N100 | -3.37 ± 3.20 | -2.11 ± 1.54 | T = -1.658, p = 0.112 |
| | P180 | 0.73 ± 5.63 | 2.30 ± 3.13 | **T = -2.764, p = 0.012**[#] |
| preSMA | N15 | -1.21 ± 9.91 | 0.13 ± 3.77 | T = -1.364, p = 0.187 |
| | P30 | 1.25 ± 7.53 | 0.41 ± 2.92 | T = 1.175, p = 0.253 |
| | N45 | -2.36 ± 2.95 | -0.76 ± 1.43 | **T = -2.330, p = 0.030**[#] |
| | P60 | 2.58 ± 3.54 | 1.08 ± 1.95 | T = 1.986, p = 0.060 |
| | N100 | -4.03 ± 3.17 | -2.34 ± 1.61 | **T = -2.620, p = 0.016**[#] |
| | P180 | 5.36 ± 6.08 | 3.01 ± 3.90 | T = 0.005, p = 3.148 |

Mean ± SD was reported for TEP peak amplitude. [#] **Bold text** indicates statistically significant results ($p < 0.05$).

Table S3. ERP components during task performance

| Target | ERP component | Trial type | Mean ± SD | Successful stop vs uncertain go | R with TEP component |
|---|---|---|---|---|---|
| M1 | P2 | successful stop | 2.24 ± 3.60 | **T = 4.418, p = 0.000**[#] | R = -0.377, p = 0.084 |
| | | uncertain go | -0.79 ± 1.16 | | R = -0.025, p = 0.913 |
| | N2 | successful stop | -3.38 ± 2.58 | T = -2.005, p = 0.057 | R = 0.350, p = 0.202 |
| | | uncertain go | -2.35 ± 1.18 | | R = 0.080, p = 0.709 |
| | P3 | successful stop | 5.79 ± 4.94 | **T = 4.099, p = 0.000**[#] | R = 0.032, p = 0.890 |
| | | uncertain go | 1.23 ± 2.22 | | R = 0.081, p = 0.734 |
| preSMA | P2 | successful stop | 2.39 ± 2.56 | **T = 6.618, p = 0.000**[#] | R = -0.162, p = 0.449 |
| | | uncertain go | -1.44 ± 1.61 | | R = 0.194, p = 0.376 |
| | N2 | successful stop | -4.66 ± 2.82 | **T = -2.242, p = 0.035**[#] | R = -0.280, p = 0.261 |
| | | uncertain go | -3.11 ± 1.86 | | R = -0.060, p = 0.791 |
| | P3 | successful stop | 6.86 ± 3.24 | **T = 7.900, p = 0.000**[#] | R = 0.310, p = 0.160 |
| | | uncertain go | 0.76 ± 1.51 | | R = 0.000, p = 1.000 |

Mean ± SD was reported for ERP peak amplitude. [#] **Bold text** indicates statistically significant results ($p < 0.05$).
